# The structural dynamics of macropinosome formation and PI3-kinase-mediated sealing revealed by lattice light sheet microscopy

**DOI:** 10.1101/2020.12.01.390195

**Authors:** Shayne E. Quinn, Lu Huang, Jason G. Kerkvliet, Joel A. Swanson, Steve Smith, Adam D. Hoppe, Robert B. Anderson, Natalie W. Thiex, Brandon L. Scott

**Affiliations:** South Dakota School of Mines and Technology (South Dakota Mines), Nanoscience and Nanoengineering, Rapid City, SD.; BioSNTR, South Dakota Mines, Rapid City, SD.; South Dakota State University (SDSU), Department of Biology and Microbiology, Brookings, SD.; BioSNTR, SDSU, Brookings, SD.; SDSU, Department of Chemistry and Biochemistry, Brookings, SD.; University of Michigan, Department of Microbiology and Immunology, Ann Arbor, MI.

## Abstract

Macropinosomes are formed by shaping actin-rich plasma membrane ruffles into large intracellular organelles in a phosphatidylinositol 3-kinase (PI3K)-coordinated manner. Here, we utilize lattice lightsheet microscopy and image visualization methods to map the three-dimensional structure and dynamics of macropinosome formation relative to PI3K activity. We show that multiple ruffling morphologies produce macropinosomes and that the majority form through collisions of adjacent PI3K-rich ruffles. By combining multiple volumetric representations of the plasma membrane structure and PI3K products, we show that PI3K activity begins early throughout the entire ruffle volume and continues to increase until peak activity concentrates at the base of the ruffle after the macropinosome closes. Additionally, areas of the plasma membrane rich in ruffling had increased PI3K activity and produced many macropinosomes of various sizes. Pharmacologic inhibition of PI3K activity had little effect on the rate and morphology of membrane ruffling, demonstrating that early production of 3’-phosphoinositides within ruffles plays a minor role in regulating their morphology. However, 3’-phosphoinositides are critical for the fusogenic activity that seals ruffles into macropinosomes. Taken together, these data indicate that local PI3K activity is amplified in ruffles and serves as a priming mechanism for closure and sealing of ruffles into macropinosomes.

## Introduction

Macropinocytosis, or “cell drinking,” is a form of clathrin-independent endocytosis that results in the non-specific uptake of large volumes of extracellular fluid and solutes. This central macrophage function enables immune surveillance, clearing of debris, and sampling of the local environment for the presence of pathogen- or damage-associated molecular patterns, cytokines, growth factors, nutrients, and other soluble cues^1–6^. Macropinosomes also serve as platforms to integrate this diverse information and to activate a variety of signaling pathways^7–10^. The major macrophage growth factor, colony-stimulating factor-1 (CSF-1), stimulates macropinocytosis and contributes to ligand-dependent modulation of CSF-1 receptor signaling^9^. Additionally, cytokines such as CXCL12, and the bacterial cell wall component lipopolysaccharide (LPS) acutely stimulate macropinocytosis^5,11,12^.

Construction of a macropinosome proceeds through autonomous, ligand-independent plasma membrane extensions known as ruffles, which are driven by actin polymerization and require the phosphorylation and dephosphorylation of different signaling phospholipids^2,13^. In the closely related process of solid particle uptake, the shape of the particle is used to template the phagosome structure^14,15^. In contrast, ruffles that fuse into macropinosomes do not have a structural framework to use as a template resulting in various potential closing mechanisms including ‘purse string’ closure of circular dorsal ruffles^13^, closure at the distal tips of ruffles^7^, and more recently described closure following actin tentpole crossing^12^. Regardless, the result is an internal organelle derived from the plasma membrane that is filled with extracellular fluid^16^.

The dynamic lipid microenvironment impacts the localization of downstream effector molecules that drives actin polymerization and ruffle growth into macropinosomes^17^. The production of 3’ phosphoinositides by PI 3-kinase (PI3K) is required to generate isolated patches of phosphatidylinositol 3,4,5,triphosphate (PIP_3_) on the plasma membrane^18,19^, and the sequential breakdown of PIP_3_ into PI(3,4)P_2_ and ultimately PI is necessary for successful macropinosome formation^20^. It is only in the cellular slime mold *Dictyostelium,* that the signal coordination throughout the 3D ruffle volume during macropinocytosis has been well described^18^, and there is still more to learn about how these events are spatially coordinated in metazoan cells^21^. The precise membrane dynamics of macropinocytosis and the spatial coordination of PI3K in forming ruffles remains unclear because of the low spatial and temporal resolution of other microscopy approaches. Previously, widefield ratiometric imaging has shown that PIP_3_ concentration peaks after ruffle circularization^17,22,23^, and PI3K inhibitors, including LY294002, have demonstrated that PI3K activity is only required for macropinosome closure, but does not inhibit ruffling^17^. Recently, high-resolution imaging of macropinocytosis in a macrophage-like cell line (RAW 264.7) indicated that the prior models of macropinocytosis may be more diverse than previously thought^12^. With the variety of different macrophage-like cell lines available, all of which perform constitutive macropinocytosis while displaying different phenotypes and protein expressions^24^, it is more important than ever to expand the 3D understanding of membrane dynamics and protein localization during macropinocytosis.

Here, we employ the powerful three-dimensional (3D) imaging capabilities of lattice lightsheet microscopy^25^ (LLSM) and volumetric image analysis to create high-resolution movies of plasma membrane dynamics and PI3K activity during ruffling and macropinocytosis. The images and movies we present advance our understanding of the spatial dynamics of membrane ruffling, the morphologies that lead to macropinosomes, the spatial distribution of PI3K activity during macropinocytosis, and finally acknowledge the structural variability among different cell lines during macropinocytosis in fetal liver (FLM), bone marrow derived (BMDM), and RAW 264.7 macrophages. Our results show that the majority of macropinosomes form by non-specific collisions of adjacent PI3K-rich ruffles. We show that PI3K activity is present at the earliest stages of ruffle extensions and is highly localized to the bottom of ruffles after the membrane has closed into a macropinosome. Finally, we modulate the rate of macropinocytosis using stimulation and pharmacological inhibition to demonstrate that the ruffle morphology is unaffected, but PI3K activity is required to prime ruffle membranes for sealing into macropinosomes.

## Results

### LLSM allows volumetric visualization of plasma membrane movements relative to PIP_3_ and PI(3,4)P_2_ distribution during macropinocytosis

Our first objective was to capture the 3D structure of the plasma membrane relative to PI3K activity during macropinosome formation. LLSM imaging was performed on fetal liver macrophages (FLMs) stably expressing the fluorescent proteins mNeonGreen localized to the plasma membrane via the lipidation signal sequence from Lck (Mem-mNG) and the pleckstrin homology domain of Akt fused to mScarlet-I (AktPH-mSc). The AktPH probe recognizes PIP_3_ and PI(3,4)P_2_ with similar affinity and has been used extensively to characterize PI3K activity during macropinocytosis^8^. Using LLSM imaging in conjunction with the molecular visualization software, ChimeraX^26^, we represent the data using the following renderings: isosurface, surface mesh, volumetric maximum intensity, and orthogonal planes. An isosurface is a three-dimensional contour map that represents points in volume space as constant values at a given intensity threshold causing all pixels above the threshold to be shown as a solid-colored voxel (value of 1), while all voxels below the threshold are transparent/hidden (value of 0) resulting in a crisp surface depiction of the membrane probe (Fig. 1a). These images were of sufficient resolution that the detailed structure of ruffles and forming macropinosomes could be observed in living cells (Fig. 1a), similar to scanning electron microscopy imaging of bone marrow derived macrophages (Fig. 1b). The isosurface is rendered with ambient occlusions meaning that any internal information is hidden from view. To visualize the recruitment of AktPH-mSc relative to the membrane, we first used volumetric intensity renderings that display the most intense color value underlying the pixel along the line of sight (ChimeraX User Guide), which provides a visually intuitive method of displaying 3D fluorescent intensities (Fig. 1c). The resulting volume is similar to a maximum intensity projection; however, the projection dimension is dependent on the orientation, i.e., top down would yield an xy-maximum intensity projection collapsed in z and a side view would provide the corresponding xz/yz maximum intensity collapsed in y/x respectively. As can be seen in the volume renderings, Mem-mNG persisted on newly formed intracellular vesicles derived from the plasma membrane (Fig. 1c). Moreover, we observed membrane movements throughout the entire formation and early trafficking of macropinosomes, as well as the recruitment of AktPH-mSc to forming macropinosomes (Fig. 1c (arrow), Supplementary Movie 1). It is difficult to perceive depth in the still-frame volumetric renderings but orthogonal plane slices (orthoplanes) in xy, yz, and xz (~0.1 μm thick) showed that AktPH-mSc was enriched in ruffles to varying degrees and intensely labeled circular structures found near the base and sides of ruffles (Fig. 1d, Supplementary Movie 2). While orthoplanes are effective for examining the 2D relationships between the fluorescent signals, they can also produce incomplete or distorted perspectives that are resolved by viewing the full volumetric data (Supplementary Figure 1), such as when a macropinosome appears closed vs open. To overcome these limitations, we implemented a mesh derived from the Mem-mNG isosurface with transparent polygonal faces (only displaying edges and vertices) that enables visualizing the underlying volumetric AktPH-mSc signals, while maintaining the structural framing needed to resolve plasma membrane rearrangements (Fig. 1e).

**Figure 1.**
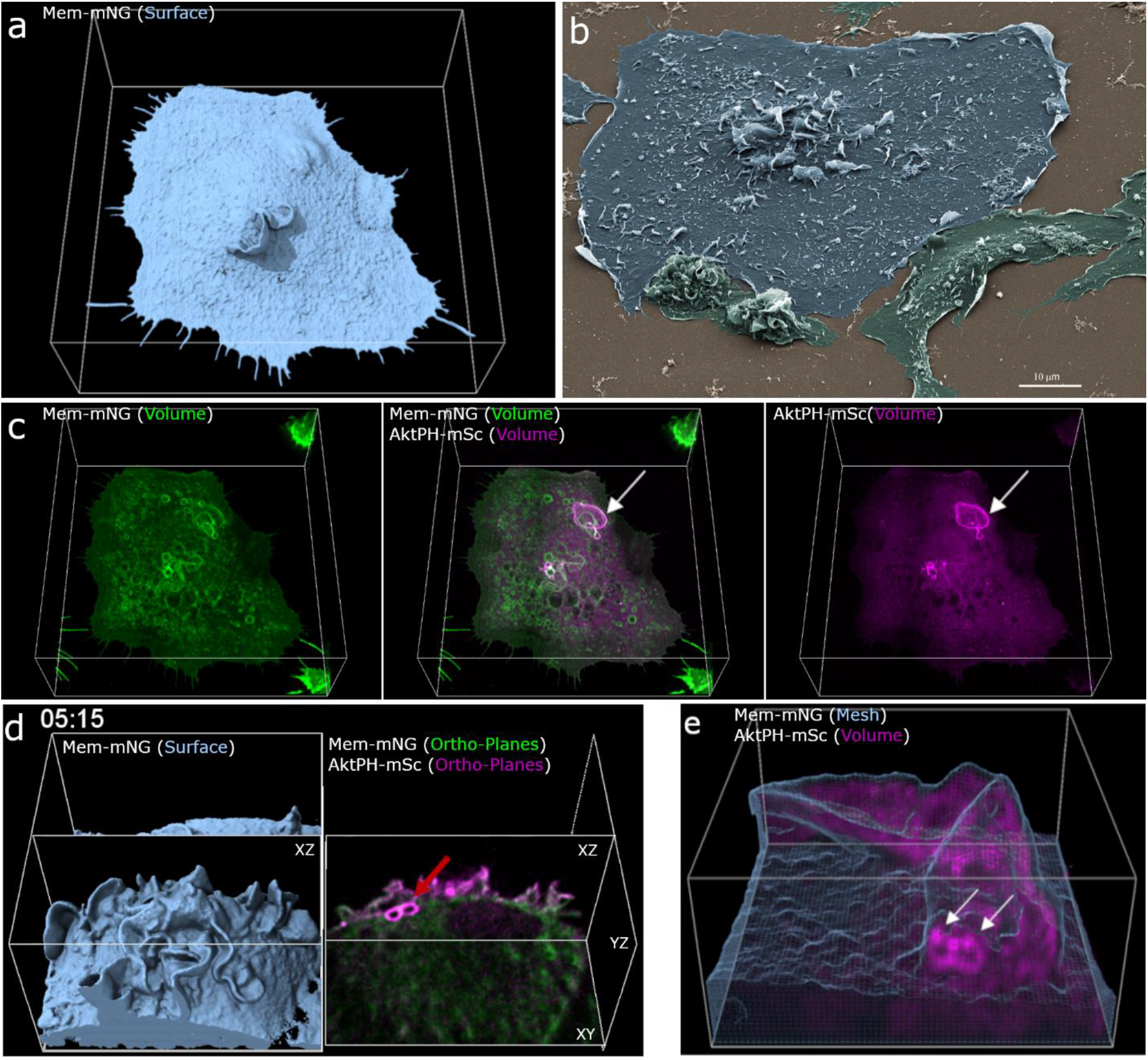
3D visualization of macrophages allows insight into membrane structure and phosphatidylinositol dynamics during macropinocytosis. a) Isosurfaces show the plasma membrane of a live cell that is actively macropinocytosing (Region 68×72×25 μm x, y, z). b) SEM image of a macrophage acutely stimulated with CSF-1 shows high-resolution fixed cells (Scale bar is 10 μm) c) Volumetric intensities show specific local fluorescence (left->right) volumetric membrane (green), dual volumetric membrane and AktPH-mSc, volumetric AktPH-mSc (magenta). Volumetric renderings provide a method to visualize the transient fluorescent intensities throughout the volume of the cell (Region 68×72×25 μm). d) Combinations of visualization techniques such as Isosurface (left) displayed alongside orthogonal planes (right) further clarify how each plane is chosen to show internal intensities (Region 29×30×19 μm). e) Mesh rendering of the Mem-mNG probe along with volumetric AktPH-mSc provides a representation of the plasma membrane structure as well as underlying fluorescence. The white arrows indicate the post closure recruitment of AktPH-mSc (Region 13×14×7 μm). Different rendering methods provide insight into cellular characteristics such as structure, depth, and fluorescent intensity and provide a foundation for visualizing localization of AktPH-mSc to the constantly changing plasma membrane during macropinocytosis.

For visualizing a macropinosome sealing event, we began by imaging a representative cell for these data that retained the canonical morphology like that of BMDMs with a profile view representing a ‘fried egg’ ranging from ~1 μm thick at the outer edge and up to ~6 μm thick near the center. Next, an XY-MIP was used to quickly find potential closures using the AktPH-mSc signal as an indicator. The identified coordinate was then translated to the corresponding isosurface, and the timepoint at which the isosurface was fully closed was considered as a nascent macropinosome. These formations were examined using the full volumetric intensity projections for further analysis. Together, combinations of these visualization techniques were applied to 122 macropinosome formations and enabled correlating the location and timing of PI3K activity to the membrane extension, curvature, and fusion of macropinosomes with unprecedented spatial and temporal resolution.

### AktPH-mSc is recruited early during ruffle expansion and peaks at the base of ruffles after macropinosome sealing

In prior analysis of macropinocytosis using microscopy methods with low axial resolution, AktPH recruitment was identified as ruffles that transitioned into closed circular ruffles and nascent macropinosomes^8^. Here, the enhanced z-axis resolution and detection sensitivity of LLSM enabled visualizing the dynamic recruitment of AktPH-mSc to ruffles as they began to protrude from the plasma membrane (Fig. 2a) until maturation where tubulation and fusion between adjacent macropinosomes occurs (Supplementary Movie 3). As these early ruffles expanded laterally along the plasma membrane and protruded vertically from the cell surface, some ruffles continued to accumulate AktPH-mSc, whereas others lost AktPH-mSc and receded back into the cell suggesting different levels of PI3K activity in neighboring ruffles influences the outcome of a ruffling region (Fig. 2b). Ruffles that maintained AktPH-mSc throughout the ruffle volume continued to grow and formed macropinosomes, which were accompanied by an intense transient recruitment of AktPH-mSc to the base of the ruffle around the nascent macropinosome (Fig. 2c, Supplementary Movie 4). Given the early localization and amplification of PI3K signaling in ruffles that become macropinosomes, we wondered in what ways PI3K activity contributed to 3D ruffle dynamics.

**Figure 2.**
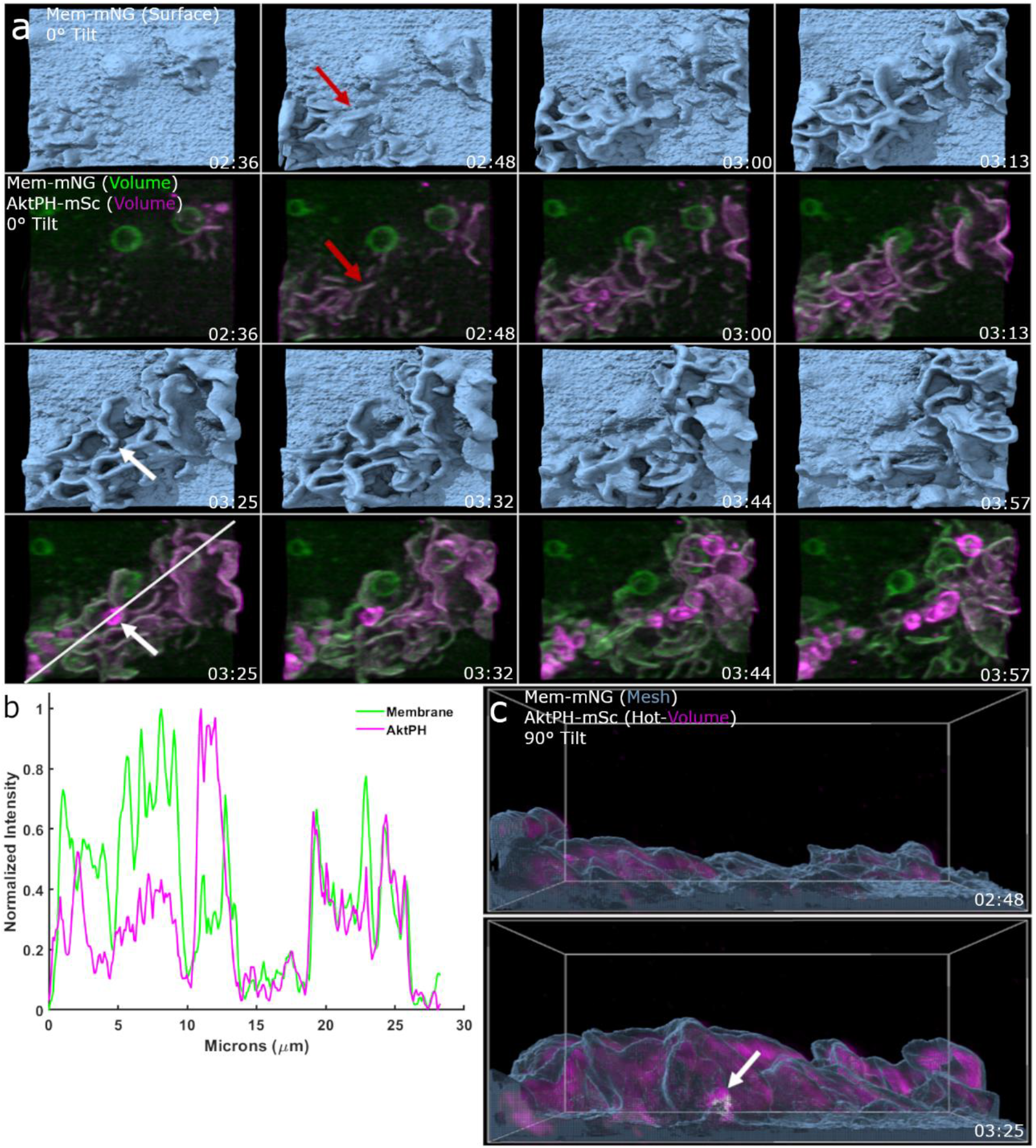
Early PI3K activity leads to amplification of PIP_3_/PIP_2_ in developing ruffles, macropinosome formation, and post closure recruitment. Top view of an Mem-mNG isosurface rendering provides depth for 3D visualization of ruffle extension. Dual-color volumetric intensity display comparing the recruitment of AktPH-mSc to early and expanding ruffles as well as sealed macropinosomes (Region 21×19um x,y). b) Intensity line-scan of the volumetric Mem-mNG and AktPH-mSc shows their co-scaled relative intensities for extending membrane ruffles, as well as recruitment around a sealed macropinosome. c) Side view of the isosurface mesh plasma membrane and volumetric AktPH-mSc (Magenta Hot color scale) from a shows that the early stages of ruffle development is filled with AktPH-mSc and the resulting macropinosome (white arrow) receives a final intense AktPH-mSc recruitment around the formed macropinosome at the bottom of the ruffle (Region 21×19×15um).

### PI3K activity is required for macropinosome sealing, but not ruffling

To gain insight into the role of PI3K in regulating the morphological dynamics of macropinocytosis, we used the broad-spectrum PI3K inhibitor LY294002, which inhibits the closure phase of macropinocytosis in macrophages^27^. Non-treated control cells formed transient dorsal ruffles that recruited AktPH-mSc and closed into macropinosomes, as seen by the surface rendering and intracellular void that is maintained in the plane view (Fig. 3a-c, Supplementary Movie 5). LY294002 treatment for 30 min did not stop ruffle formation, but eliminated AktPH-mSc recruitment to membrane ruffles (Fig. 3d,e, Supplementary Movie 6). Furthermore, LY294002-treated cells formed ruffles that appeared to close into a macropinosome but retracted back to the plasma membrane and failed to maintain an intracellular organelle (Fig. 3f). In order to further understand the dynamic cellular response to PI3K inhibition, we imaged the same cells immediately before and after LY294002 treatment. Immediately after treatment, PI3K activity was halted as evidenced by clearance of AktPH-mSc from the PM ruffles and diffuse cytosolic localization (Supplementary Figure 2). Interestingly, many of the ruffles receded into the plasma membrane, but ruffling resumed within 10 min and was similar to the untreated cells by 30 min (Supplementary Movie 7). Taken together, these data suggest that PI3K activity is dispensable for ruffle formation but is required for membrane sealing to generate a macropinosome.

**Figure 3.**
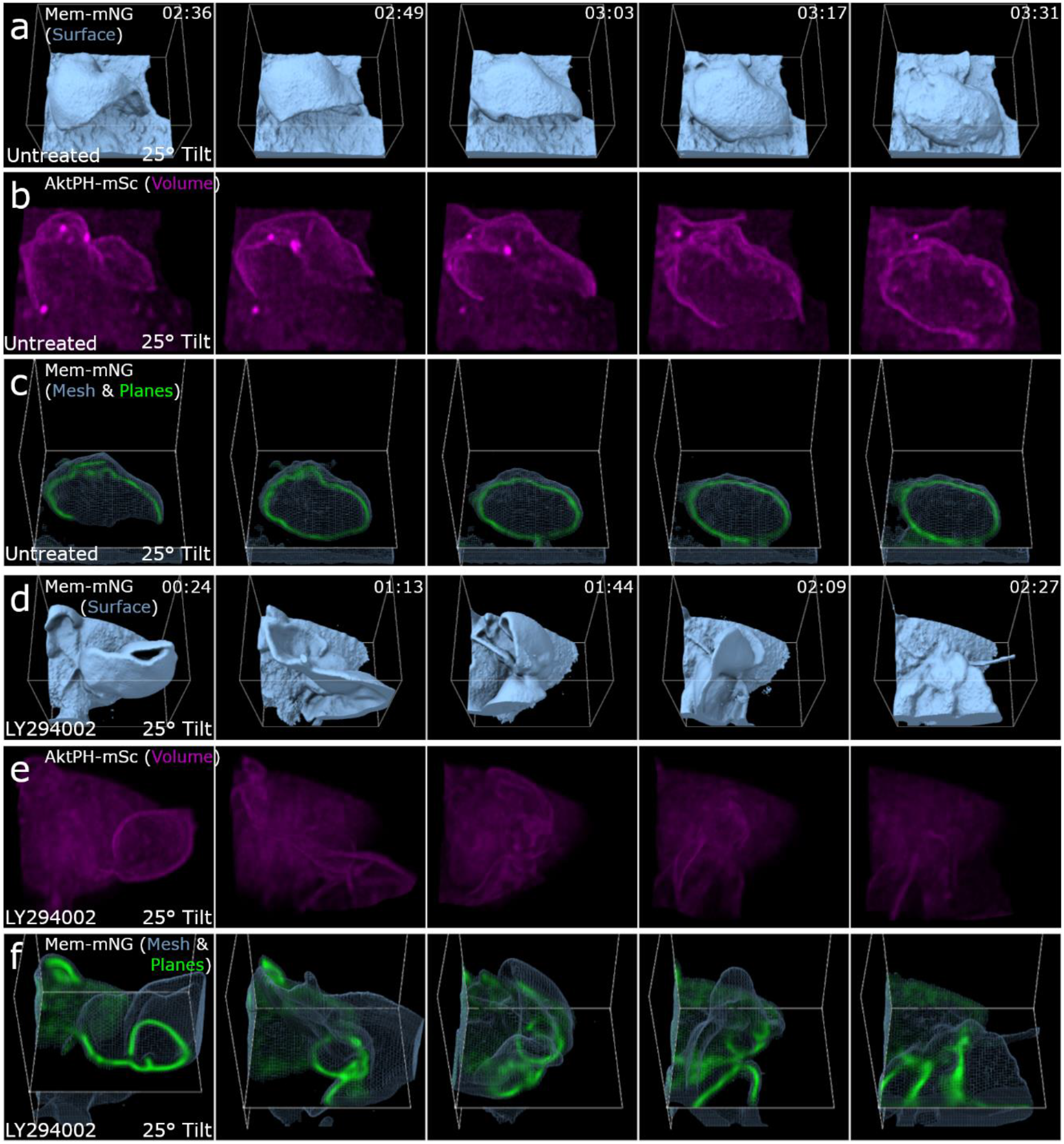
PI3K activity is required for membrane sealing and separation from PM/internalization of a complete macropinosome but not membrane ruffling. a) Isosurface rendering of Mem-mNG for an untreated macrophage during a successful macropinocytosis event where the sheet curls back toward the membrane for fusion/sealing (Region 12×13×10 μm). b) Volumetric rendering of Sc-AktPH of the untreated cell shows the increase of PI3K activity in the ruffle that creates a macropinosome (Region 12×13×10 μm). c) Mesh and orthogonal planes of Mem-mNG show the internal membrane organization of the ruffle and resulting macropinosome (Region 12×13×10 μm). d) Isosurface rendering of an LY294002 treated macrophage provides depth to the attempted closure of a macropinocytic cup (Region 10×12×10 μm). e) Volumetric intensity rendering of Sc-AktPH for an LY294002-treated macrophage shows the diffuse distribution of AktPH and minimal PI3K activity. The cytosolic intensities were co-scaled for the untreated and treated macrophage (Region 10×12×10 μm). f) XY-plane for the Mem-mNG probe of an LY294002-treated cell during a failed macropinocytosis event. In the surface view, the ruffle appeared to form a macropinosome; however, when overlaid with the plane view is became clear that it failed to fully form into a macropinosome. The ruffle quickly reduced in size and became undistinguishable within the cytosol, while never receiving the post closure increase of PI3K activity (Region 10×12×10 μm).

### PI3K activity primes ruffles for fusion to seal into nascent macropinosomes

We next sought to categorize the macropinosome formation based on the way the membrane fused and the relative amount of PI3K activity. Based on the previously established models for macropinosome formation there would be distinguishing characteristics depending on the sealing formation method. Either we would find linear extensions where the distal tips would collide, circularize, and seal, or we would find filopodial spikes that form and the membrane would fill in the space between before twisting to seal^12^. Surprisingly, we found that approximately 88% of the quantified macropinosomes formed when the leading edge of extending ruffles collided along the sides of nearby membrane surface or ruffles that typically only involved a small portion of the second ruffle, so long as the ruffle area had elevated AktPH-mSc (Fig. 4). The remaining 12% of events we observed were classified as tidal-wave like structures in which a mostly isolated planar ruffle extended from the cell surface where the entire ruffle was rich in AktPH-mSc, the ruffle gained curvature in a rolling fashion, and resulted in fusing back with the plasma membrane rather than an extending ruffle (Supplementary Movie 5). However, given that the entire ruffle area was rich in AktPH-mSc, these types of formation follow the same underlying mechanism as collisions with adjacent membrane extensions. Frequently, a single ruffle area produced multiple macropinosomes and were the result of similar but smaller ruffle extensions that quickly fused near the base of larger ruffles (Fig. 4). Within the ruffle, forming macropinosomes recruited AktPH-mSc near the base of the ruffle as they transitioned into a spherical shape prior to detaching from the plasma membrane and moving independently (Fig. 4, Supplementary Movies 3,8). We hypothesized that regions with highly concentrated AktPH-mSc localization would correlate with increased macropinocytic activity. Indeed, this phenomenon was observed in four of the eleven constitutive cells (Fig. 5). These ruffling regions resulted in the formation of many macropinosomes through the intersection of multiple ruffles that were nearly indistinguishable from one another and only became apparent through the PI3K post closure activity (Fig. 5d,e). Therefore, the elevated PI3K activity creates a microenvironment suited for the rapid fusion of PI3K-primed ruffles into macropinosomes of various sizes within short timeframes (Supplementary Movies 9,10). We hypothesized that other signaling that activates PI3K activity may stimulate distinct ruffling morphologies or modulate the rate of macropinocytosis.

**Figure 4.**
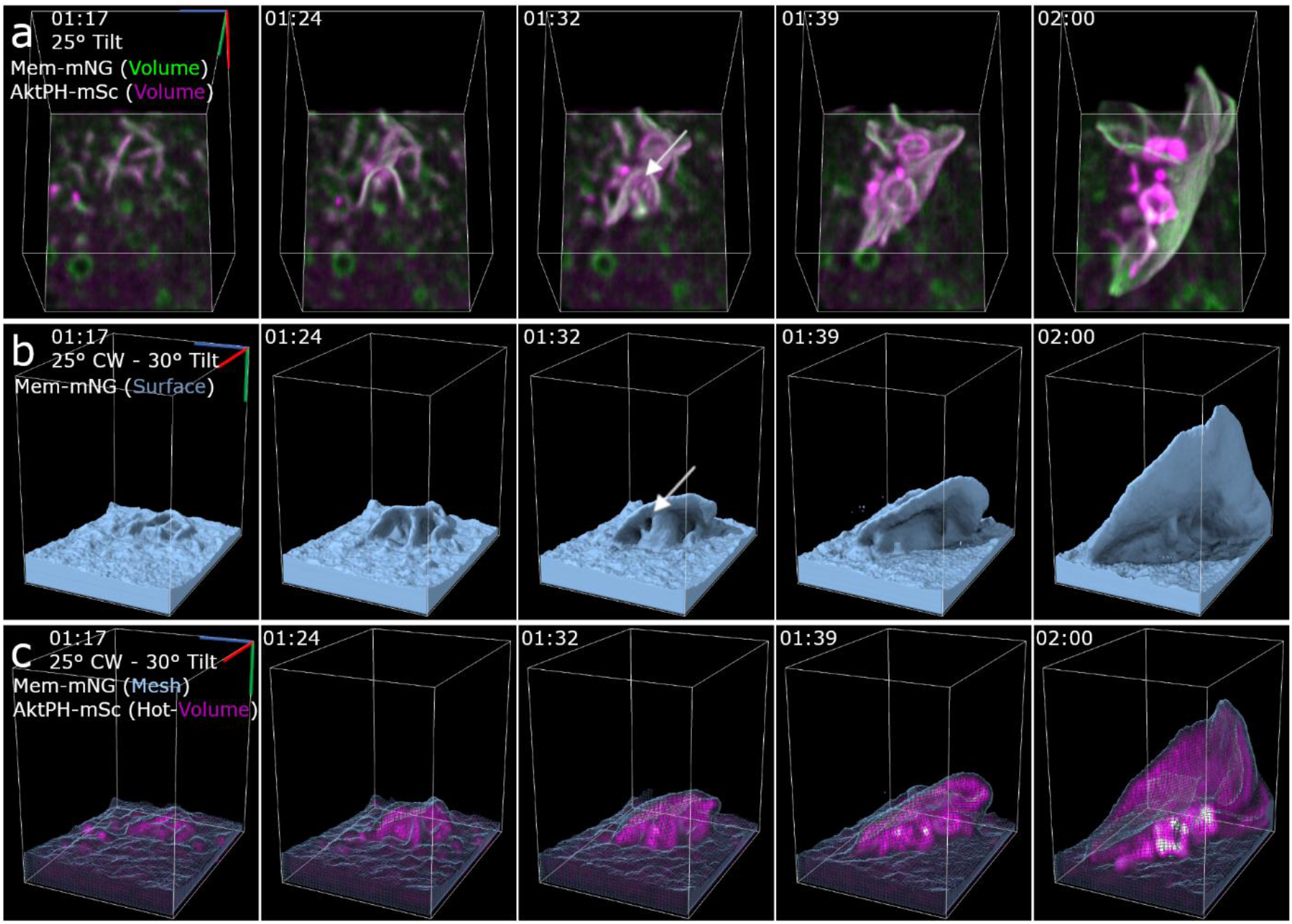
Macropinosomes form via PI3K-primed ruffle fusion. a) Dual volumetric intensities of Mem-mNG and AktPH-mSc show the intensity of each probe as the ruffles and macropinosomes form. The montage shows the earliest stage of the ruffle that extends vertically and forms macropinosomes along the length near the base of the primary ruffle as a result of smaller AktPH-mSc-rich extensions colliding. The white arrow points at the macropinosome forming region further emphasized in the isosurface (Region 9×12×13 μm). b) Isosurface rendering of Mem-mNG shows the structure of the extending ruffle and the continued sheet extension after the macropinosomes formed. The white arrow emphasizes the small pocket that closes to form one of the macropinosomes (Region 9×12×13 μm). c) Mesh surface rendering of Mem-mNG and volumetric AktPH-mSc shows the internalized macropinosome with the increased localization of AktPH-mSc at the bottom of the ruffle (Region 11×9×12 μm).

**Figure 5.**
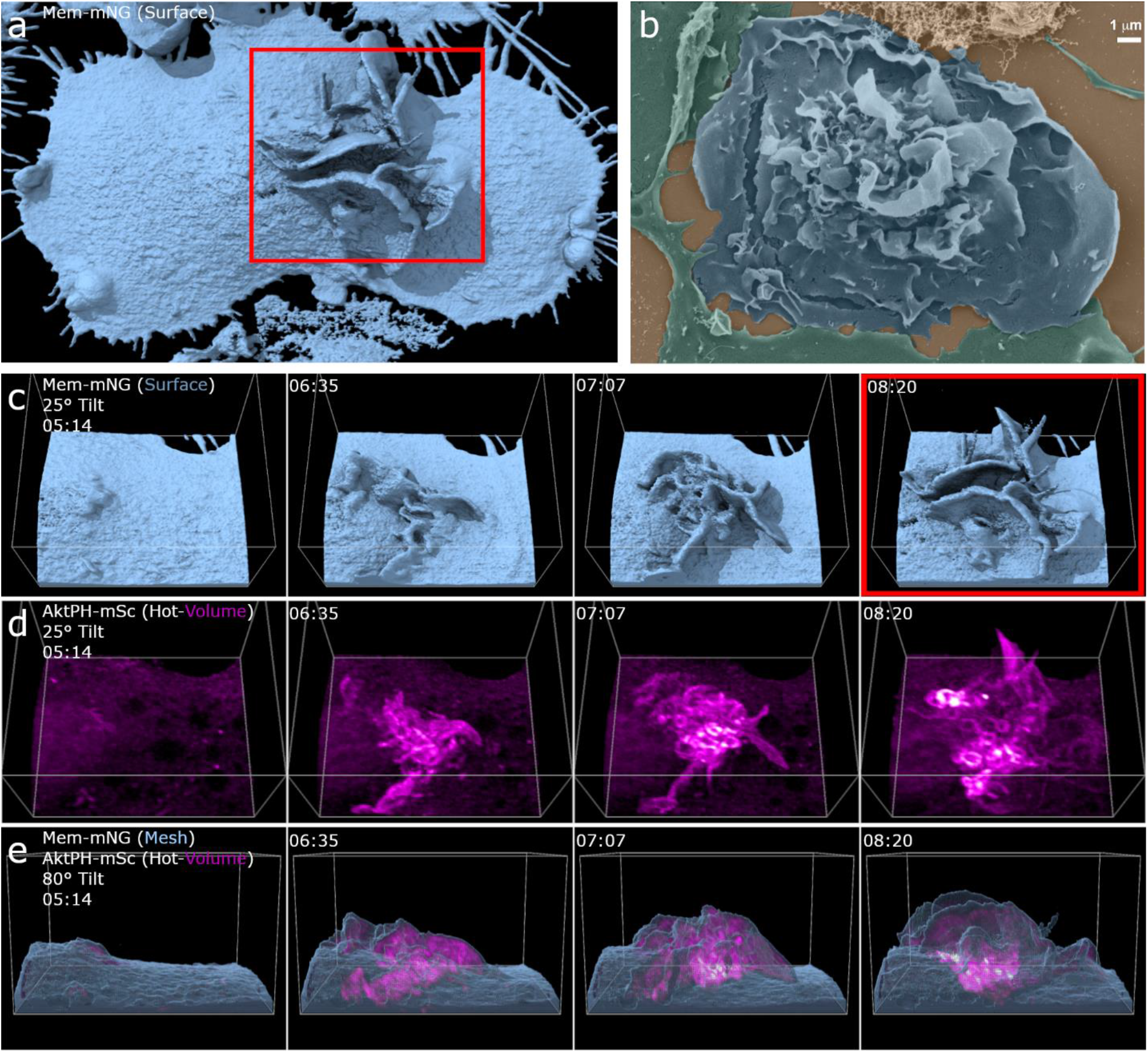
Phosphatidylinositol localization and chaotic ruffling underlie macropinocytosis in complex membrane structures. a) Single time point, full cell surface rendering of chaotic macropinocytosis event. The red box correlates to the same frame in c-d. b) SEM images of a BMDM showing similar highly active ruffling regions. Scale bar 1 μm c) Isosurface montage shows the chaotic orientation of membrane structure (Region 27×22×16 μm, 25° tilt). d) Volumetric AktPH-mSc (Magenta-Hot) provides a more detailed emphasis on the AktPH activity within the membrane ruffles and highlights the macropinosomes that have formed (Region 27×22×16 μm, 25° tilted). e) Mesh Surface with AktPH-mSc (Magenta Hot) shows the AktPH activity as the ruffle develops as well as the increased recruitment around formed macropinosomes at the base of the event.

### CSF-1 growth factor signaling promotes extensive circular ruffling and macropinocytosis

CSF-1 is an essential macrophage growth factor that stimulates macropinocytosis at levels controlled by the concentration of the CSF-1 signal^9^. Macrophages starved of CSF-1 for 24 hrs and then acutely stimulated produced expansive circular ruffles that initiated from the distal cellular margins coincident with cellular spreading (Fig. 6). LLSM imaging revealed a circular ruffle that initiated at the edge of the cell with a height of approximately 2 μm above the dorsal surface and constricted to a central location in a coordinated manner (Fig. 6a). A striking feature of this ruffle was the confinement of AktPH-mSc within the limiting edge of the ruffle. As the circular ruffle constricted toward the center of the cell, AktPH-mSc was highly concentrated within it and was nearly undetectable in the rest of the cell (Fig. 6b, Supplementary Movie 11). In the volumetric projections, many macropinosomes were observed to form during the constriction process without additional membrane protrusions being generated. This is consistent with our hypothesis that increased PI3K activity primes membranes for fusion to generate macropinosomes (Fig. 6c, Supplementary Movies 11,12). Thus, CSF-1 initiated long range signaling and PI3K activation resulting in coordinated movements of the cytoskeleton throughout the cell.

**Figure 6.**
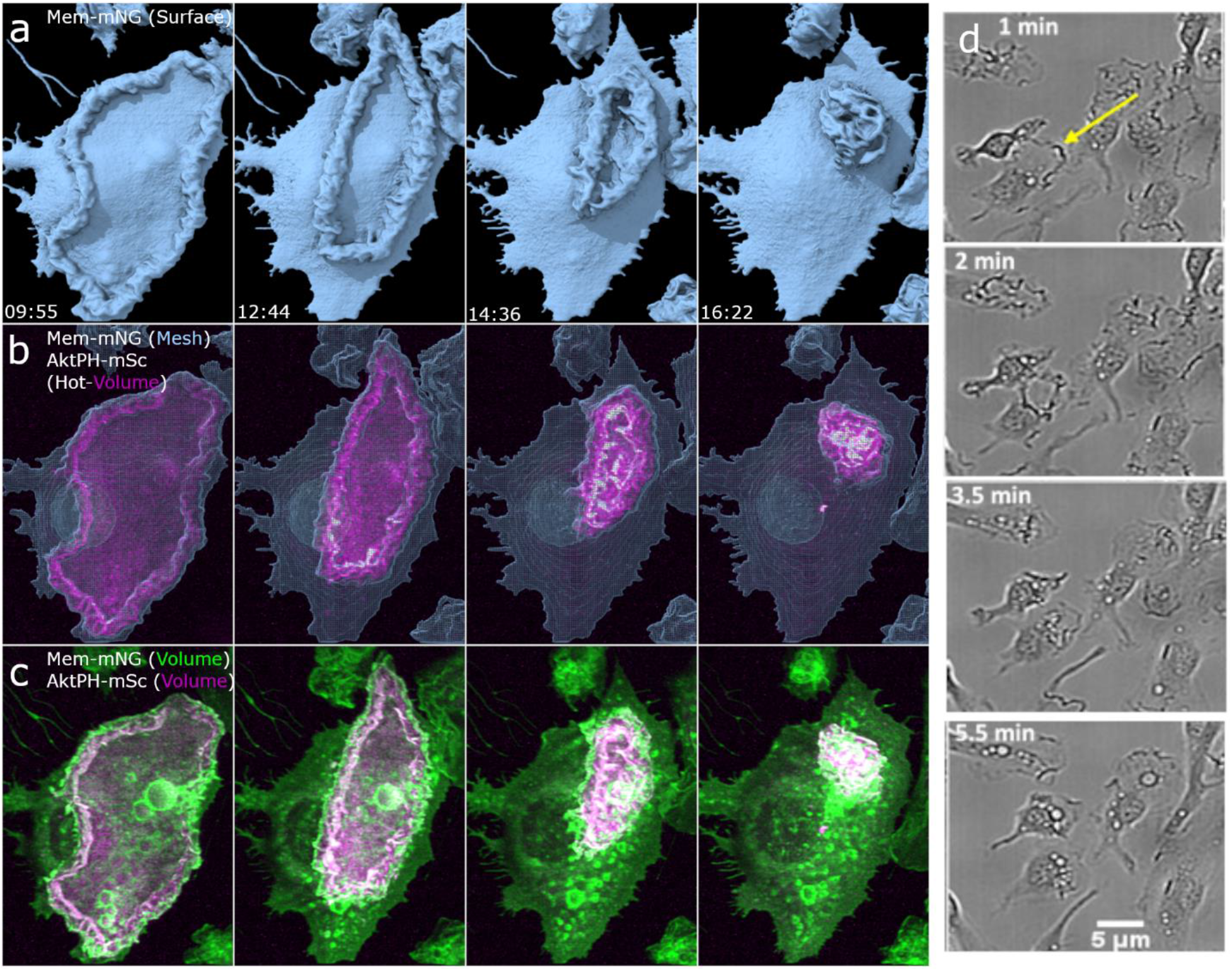
Growth factor starvation and stimulation results in the formation of large circular dorsal ruffles that corrals PIP_3_/PIP_2_. Macrophages were starved of CSF-1 for 24 h, imaged for 5 minutes as a baseline, and imaging restarted 1 min after stimulation with CSF-1. Four-frame montages provide a visual display of the large dorsal ruffle that acts as a diffusional barrier that restricts PIP_3_/PIP_2_ to the inside of the ruffle as it is cleared from the surface. This barrier is likely acting as a signal amplification mechanism stimulating the production of many macropinosomes. a) Isosurface rendering provides crisp surface directionality, b) Surface mesh and volumetric AktPH-mSc (magenta-hot), show the restricted probe as the membrane converges c) Volumetric Intensity of both Mem-mNG and AktPH-mSc show the intensity locations of the membrane ruffle and the restricted AktPH (49×60 μm x,y). d) Bright field images showing multiple cells responding to stimulation with similar dorsal membrane clearing.

### LPS stimulates regional ruffling and generates large numbers of macropinosomes

The bacterial cell wall component lipopolysaccharide (LPS) activates PI3K through the Akt pathway^28^, and acutely stimulates macropinocytosis^3^. Recently, LPS stimulation was used to characterize a novel formation mechanism involving actin tentpoles supporting membrane veils which cross to create a macropinosome^12^. When FLMs were exposed to LPS, regional patches of membrane ruffling were generated that migrated around the dorsal surface of the macrophage (Fig. 7a, Supplementary Movie 13) in a manner distinct from the dorsal surface ruffle generated by CSF-1 stimulated cells (Fig. 6); however, this process was similar in appearance to constitutive macropinocytosis (Fig. 7c, Supplementary Movie 14). The patches of ruffles in LPS-treated cells generated many small ruffles, had elevated PI3K activity and were more efficient at forming macropinosomes as compared to control (Fig 7d). Thus, the nature of macropinosome formation is coordinated over different length scales with differing intensities depending on the nature of the activating stimulus. Regardless, PI3K activity delineates ruffles and regions of the plasma membrane where macropinosomes form.

**Figure 7.**
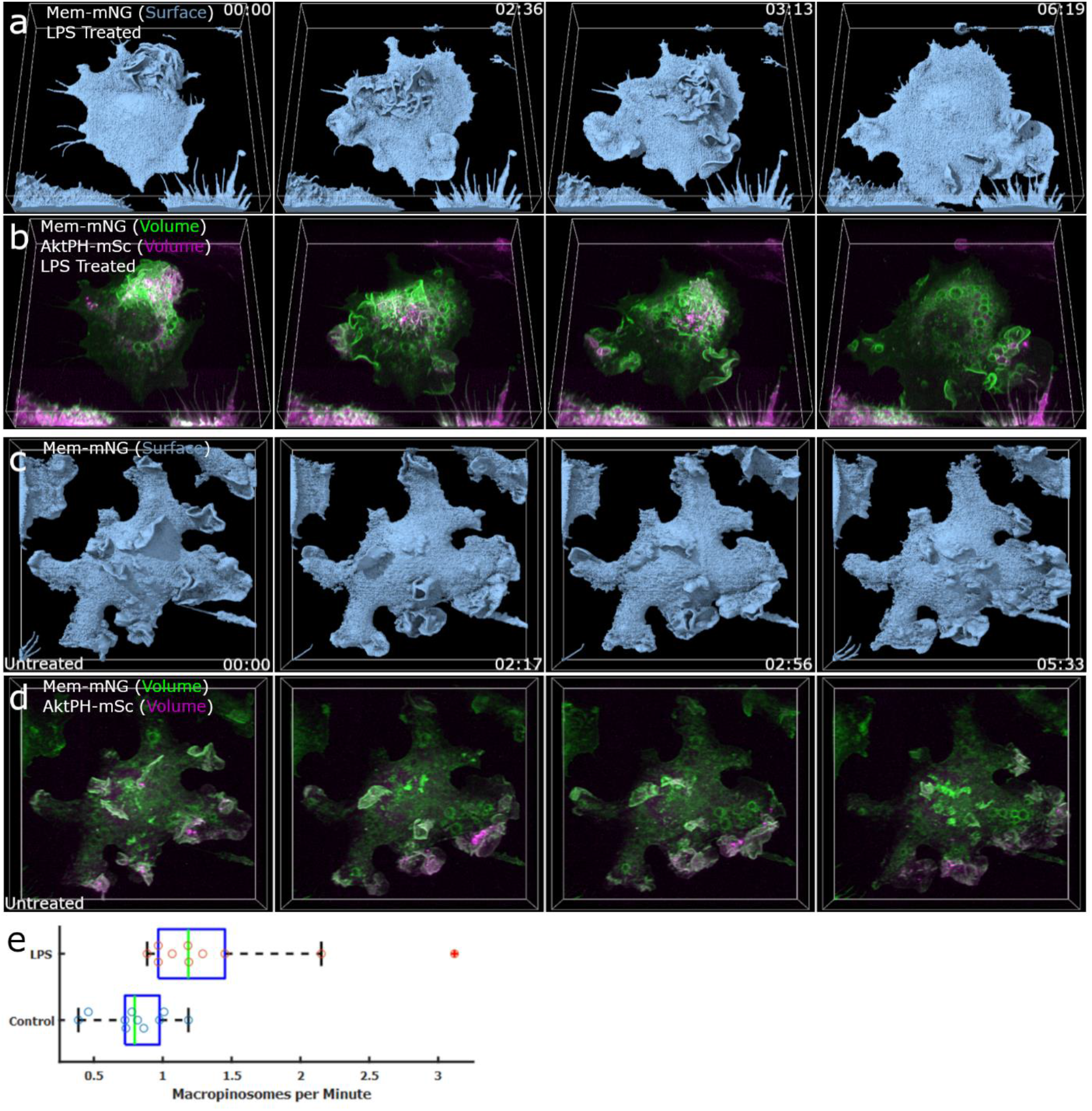
LPS stimulation increases membrane ruffling and macropinocytosis. Macrophages were pretreated with 100 ng·ml-1 LPS for 24 h prior to imaging. a) Surface rendering of Mem-mNG on an LPS stimulated macrophage provides a surface level understanding of the membrane, exploration, ruffling, and PM structure (Region 68×69×16 μm). b) Dual-color volumetric intensity projections of Mem-mNG and AktPH-mSc for an LPS stimulated cell provided the intensity activity during increased macrophage activity and shows the highly AktPH rich regions of membrane ruffling (Region 68×69×16 μm). c) Untreated macrophage Isosurface displaying less exploratory behavior (Region 68×64×16 μm). d) Dual-color volumetric intensity rendering of the untreated macrophage gives insight on the AktPH activity inside of the cell during macropinocytosis and allows for the quantitative comparison of macropinosomes formed between the stimulated and unstimulated cells (Region 68×64×16 μm). e) Box plot showing the difference in macropinocytic activity between untreated and LPS treated macrophages. All macropinosomes greater than 1 μm were manually counted using a z-projection MIP in Fiji and were distinguished by the post closure spike in AktPH-mSc intensity.

## Discussion & Conclusion

Here, we have utilized lattice light sheet microscopy to develop a new level understanding of the structural dynamics and PI3K signaling underpinning macropinocytosis. Until recently, dynamic processes such as macropinocytosis were characterized using optical techniques with poor axial resolution and elevated phototoxicity leading to subsampling the spatial and temporal dynamics and requiring inference from multiple methods such as scanning electron microscopy of fixed cells to address the formation mechanism of macropinosomes. Lightsheet microscopy overcomes these obstacles and enables us to record, with sufficient spatial and temporal resolution, the complete evolution of membrane ruffles and the mechanism by which these ruffles form into macropinosomes while also measuring the redistribution of signaling molecules controlling these processes. We have shown that macropinosomes form through several possible morphologies; however, in each case PI3K activity primes ruffles for fusion with adjacent primed membranes to form macropinosomes. In areas with elevated PI3K activity, either naturally or through external stimulation, there was an increased ruffle density leading to an increased probability of primed ruffles colliding to form macropinosomes. This model of macropinosome formation relying on PI3K priming rather than a defined geometry also explains the variation in diameter that is a hallmark of macropinosomes. The improved sensitivity of LLSM enabled detection of PI3K activity at the earliest stages of ruffle development that grows in curving ruffles and peaks around macropinosomes post closure. Our data are consistent with a mechanism driven by the geometry of curving ruffles that confines PI3K, thereby amplifying the signal, which in turn activates yet unknown fusogenic protein(s) localized to the ruffle edges mediating sealing during membrane collisions. This conclusion is supported by the observations that inhibition of PI3K activity with LY29004 did not substantially alter membrane ruffling structure, curvature, or collisions but completely inhibited sealing, even when fully spherical morphologies were observed that then collapsed back into the cell surface. These data are consistent with previous work using LY294002 as a broad-spectrum PI3K inhibitor and the extensive work done on PI3-kinase in dictyostelium^29,30^

The model suggested in this work contrasts with a recent report in LPS-activated RAW 264.7 cells that described F-actin-rich filopodial “tentpoles” protruding from the surface that twisted to constrict veils of membrane that then became macropinosomes^12^. Using Mem-mNG and Lifeact-mSc in RAW 264.7 cells under identical imaging conditions, we observed filapodial extensions that fit with the tentpole formations described previously (Supplementary Figure 3). However, we also transduced BMDM with the same NG-membrane probe and found no tentpole like structures; rather the FLM and BMDM were similar in their flat and smooth dorsal surfaces with large lamellar sheet-like ruffles (Supplementary Figure 3), suggesting that the tentpole-like formation is dispensable for macropinocytosis and may be unique to the RAW264.7 cell line.

Our data demonstrate that macropinosomes form from a variety of morphologies in macrophages, contrary to the morphologies in other cells such as the tightly controlled process observed in Dictyostelium. Specifically, macropinocytosis in Dictyostelium appears to involve the formation of a cup with a clearly segregated PIP3 containing domain^29^ whereas, our observations as well as the work of Condon et al using LLSM indicates that initiation of macropinosomes by macrophages arises from a morphologically diverse and somewhat chaotic ruffling process. Importantly, the well-defined PIP3 containing domains observed in Dictyostelium appear similar to the macropinosomes in macrophages at the sealing stage, in which substantial PIP3 products accumulate (Fig. 2). We speculate that these differences are attributed to the importance of macropinocytosis in *Dictyostelium* as a method of nutrient gathering^30,31^, while macrophages perform constitutive macropinocytosis in order to sample the environment for antigen presentation^2,7^.

Taken together, our experiments indicate a mechanism for macropinosome formation requiring amplified PI3K signaling within ruffles that become macropinosomes and that PI3K contributes primarily to priming membranes for sealing. The membrane probe and visualizations we have described set the foundation to enable rigorous testing of this mechanism using specific inhibitors and sensors that bind to the various products to determine how each step is regulated during macropinocytosis.

## Methods and Materials

### Plasmids

pCMV-VSV-G was a gift from Bob Weinberg (Addgene plasmid #8454; http://n2t.net/addgene:8454; RRID:Addgene_8454)^32^. psPAX2 was a gift from Didier Trono (Addgene plasmid #12260; http://n2t.net/ addgene:12260; RRID:Addgene_12260). pLJM1-EGFP was a gift from David Sabatini (Addgene plasmid #19319 ; http://n2t.net/addgene:19319 ; RRID:Addgene_19319)^33^. Lck-mScarlet-I and Lifeact-mScarlet-I were gifts from Dorus Gadella (Addgene plasmids #98821, #85056; http://n2t.net/addgene:98821; http://n2t.net/addgene:85056; RRID:Addgene_98821, RRID:Addgene_85056)^34^.

### Construction of the membrane and AktPH probes

The membrane probe was constructed by combining the membrane localization motif (MGCVCSSNPE) from Lck^34^ in frame with mNeonGreen in the pLJM1 backbone containing the puromycin resistance gene. The AktPH-mSc probe was constructed by using the pleckstrin homology domain from Akt in frame with mScarlet-I in the pLJM1 backbone, modified to contain the blasticidin resistance gene. Sequences were codon optimized for expression in mouse cells, synthesized, and sequence verified by GenScript (Piscataway, NJ).

### Viral transduction

Sequence-verified plasmids containing genes encoding FP-chimeras, plus the packaging plasmids pVSV-G and psPAX2 were transfected into NIH 293T cells for packaging using linear 25 kDa polyethylenimine (PEI) as a transfection reagent. NIH 293T-cell culture supernatant containing lentiviral particles was collected and added to either BMDMs, RAW 264.7, or FLMs each treated with cyclosporin A (10 μM) for two days. Transduced FLMs were selected with puromycin and blasticidin (10 μg·ml^−1^ each).

### Macrophage culture media

DMEM/F-12 (Gibco) supplemented with 20% Heat-inactivated FBS (R&D Systems), 1% Penicillin/Streptomycin (Gibco), 50 ng·ml^−1^ mCSF-1 (BioLegend), and 5 μg·ml^−1^ plasmocin (Invivogen) maintained at 37°C with 7.5% CO_2_.

### PIP_3_ inhibition (LY294002)

Coverslips were prepared as described in ‘Cell culturing and coverslip plating’ and moved to a well of media containing 50 μM LY294002 where it was incubated for 30 minutes at 37°C and 7.5% CO_2_ prior to imaging. The coverslip was then transferred to the LLSM bath containing Imaging Media, 1.7 mM Glucose, and 50 μM LY294002. The coverslip was explored using the LLS software, and three cells were chosen per coverslip that provided the best visual representation of the population. Each cell was imaged under the same parameters as described in ‘Lattice light sheet microscopy imaging’. Additionally, coverslips were tested to ensure the 30 min treatment was sufficient by imaging a coverslip, introducing LY294002 at 50 μM, and imaged three cells representing the full 30 minute treatment. Finally, untreated coverslips were placed on the microscope and three cells were selected. The three cells were imaged prior to drug treatment in the imaging bath then LY294002 was introduced at 50 μM and each cell was reimaged in reverse order (1-2-3-3-2-1). Alongside these tests a DMSO control was performed to further ensure the proper treatment.

### CSF-1 stimulation

The coverslip was starved overnight in DMEM/F-12 with 10% FBS and 1% pen/strep. After 24 h the coverslip was moved to the LLS bath containing 7 mL L-15 imaging media, and 1.7mM Glucose. The coverslip was explored using the LLS software and three cell targets were chosen and imaged for a pre-stimulation comparison. Immediately after imaging the third cell, CSF-1 was introduced at 50 ng ·mL^−1^ to the 7 mL bath. The third cell was once again imaged < 1 min after stimulation and each additional cell was imaged in reverse order (Imaging order 1-2-3-3-2-1).

### Lipopolysaccharide stimulation

FLMs were stimulated with 100 ng·ml^−1^ Lipopolysaccharides from *Salmonella enterica* serotype enteritidis (Sigma) for 24 h in culture media before being transferred to imaging media containing 100 ng·ml^−1^ for LLSM experiments.

### Macrophage isolation

Fetal liver macrophage (FLM) cell cultures were generated as described previously^35,36^. Livers were isolated from gestational day 15-19 mouse fetuses from C57BL/6J mice (The Jackson Laboratory, Bar Harbor, ME) in accordance with South Dakota State University Institutional Animal Use and Care Committee. Liver tissue was mechanically dissociated using sterile fine-pointed forceps and a single-cell suspension was created by passing the tissue through a 1 ml pipette tip^35^. Cells were plated on non-tissue culture treated dishes and kept in growth and differentiation medium containing the following: 20% heat-inactivated fetal bovine serum; 30% L-cell supernatant, a source of M-CSF^37,38^; and 50% Dulbecco’s modified growth medium containing 4.5 g·L^−1^ glucose, 110 mg L^−1^ sodium pyruvate, 584 mg·L^−1^ L-glutamine, 1 IU ml^−1^ penicillin and 100 μg·ml^−1^ streptomycin. FLM were cultured for at least 8 weeks prior to transduction and experiments.

### Cell Culturing and Coverslip Plating

FLMs were cultured in T-25 tissue culture flasks using the following culture media: DMEM F-12 with 20% FBS, 1% penicillin/streptomycin, 50 ng·ml^−1^ CSF-1, and 5 ng·ml^−1^ plasmocin. The cell cultures were split at ~85% confluence, first by washing the T-25 flask with 3 mL of PBS (-Ca/-Mg) 2 times. The cells were then lifted from the T-25 flask using 4 mL of 4°C PBS (-Ca/-Mg, +0.98mM EDTA) with gentle pipet washing for approximately 10 min. The lifted cells were moved to a 15 mL centrifuge tube (1 mL of culturing media was added to 15 mL tube if cells took >10 min to lift) and centrifuged at 200 x g for 5min. While the cells were being spun down, the T-25 flask was washed with PBS (-Ca/-Mg), filled with 5mL of culture media, and placed in the 37°C incubator to reach appropriate culture conditions. Once the cells were finished being spun down, the supernatant was removed from the 15 mL tube and the cells were resuspended in 1 mL culturing media and counted. The counting was done by mixing 10 μL of suspended cells with 10 μL of trypan blue and placed on a glass slip to be counted using a countess. The FLMs were re-plated in the original T-25 flask with ~6-7 × 10^5^ cells. The cells were washed every 2 days and given fresh media until reaching ~85% confluence where they would then be split. Cell lines were kept for approximately 2 months before being replaced with early state frozen aliquots. Macrophages were prepared for LLS imaging 24 h prior to imaging using 5mm glass coverslips. The coverslips were soaked in 90-100% ethanol and flame cleaned using a butane flame. Approximately 5 flame cleaned coverslips were placed per well of a 12 well plate each containing 1 mL of culture media. Cells were added to each well during the cell culture process described above at ~3×10^5^ cells to each 3.5cm^2^ (12-well plate) for imaging. The FLMs incubated on the flame cleaned glass coverslips in culturing media for 24 h prior to imaging. The coverslips were transferred to the LLSM bath that was filled with 7 mL of Leibovitz’s L-15 Media (supplemented with 1.7 mM glucose) at ~37°C.

The RAW 264.7 macrophages were purchased from ATCC and maintained using the suggested subculture routine. Briefly, cells were split between 70-80% confluence, reseeded at 2-4 × 10^4^ cells cm^−2^, and cultured in DMEM F-12, requiring 7.5% CO_2_ at 37°C. Due to the variability in adherence, cells were lifted using the recommended cell scraper protocol. The same protocol as discussed above was used for plating cells for imaging on the LLSM.

### Lattice Light Sheet Microscopy

The LLSM is a replica of the design described by Chen et al.^25^, built under license from HHMI. Volumetric image stacks were generated using dithered square virtual lattices (Outer NA 0.55, Inner NA 0.50, approximately 30 μm long) and stage scanning with 0.5 μm step sizes, resulting in 254 nm deskewed z-steps. Excitation laser powers used were 18μW (488 nm) and 22μW (561 nm), measured at the back aperture of the excitation objective. The emission filter cube (DFM1, Thorlabs) comprised a quadband notch filter NF03-405/488/561/635 (Semrock), longpass dichroic mirror Di02-R561 (Semrock), shortpass filter 550SP (Omega) on the reflected path, and longpass filter BLP01-561R (Semrock) on the transmitted path; the resulting fluorescence was imaged onto a pair of ORCA-Flash4.0 v2 sCMOS cameras (Hamamatsu). The camera on the reflected image path was mounted on a manual x-y-z translation stage (Newport 462-XYZ stage, Thorlabs), and the images were registered using 0.1 μm Fluoresbrite YG microspheres (Polysciences). The image capturing rates varied between 5-10 seconds per volume using 8-12 ms planar exposures depending on the brightness of the cell and imaging region.

### Scanning Electron Microscopy

BMDM were plated onto 13 mm diameter, circular glass coverslips and cultured overnight in RPMI with 10% FBS (R10). To stimulate macropinocytosis, BMDM were incubated 30 min in phosphate-buffered saline (PBS), then 15 min in PBS containing 10 nM CSF-1. Cells were fixed in 2% glutaraldehyde, 0.1 M cacodylate buffer, pH 7.4, 6.8% sucrose, 60 min at 37 °C. Fixative was replaced with a second fixative consisting of 1% OsO_4_ in 0.1 M cacodylate buffer, pH 7.4, for 60 min at 23 °C. The second fixative was replaced with 1% tannic acid in cacodylate buffer, 30 min at 4 °C, the rinsed with 3 changes of 0.1 M cacodylate buffer. Coverslips were transferred through successive changes of acetone-water mixtures, progressively increasing acetone concentrations to 100% before a final incubation in hydroxymethylsilazidane (HMDS, EM Sciences). HMDS was removed and coverslips were dried for 2 days. Coverslips were shadowed with gold and observed on a Amray 1900 field emission scanning electron microscope.

### Deconvolution and Post Processing

The raw volumes acquired from the LLSM followed the standard protocol including deskewing, deconvolving and rotating to coverslip coordinates in LLSpy^39^ as the raw data acquired from LLSM is captured at an approximate angle of ~32° with respect to the coverslip. We applied a fixed background subtraction based on an average dark current image, 10 iterations of Lucy-Richardson deconvolution with experimentally measured point spread functions for each excitation followed by rotation to coverslip coordinates and cropping to the region of interest surrounding the volume for visualization. We optimized the illumination for minimal photobleaching to occurred during the experiments. The fully processed data was opened as a volume map series in UCSF ChimeraX and utilized isosurface, mesh, 3D volumetric intensities, and orthogonal planes renderings to exam the data. The surface and mesh options utilize a three-dimensional analog of an isoline called an isosurface that represents points in volume space as constant values which were used to display the membrane probe. The isosurface provides a defined surface for the membrane probe resulting in shadowing providing visual depth to the three-dimensional data. The mesh rendering offers a similar surface definition while also providing an option to include the internal fluorescent AktPH-mSc signal. We used a volumetric intensity projection to visually display the localization of AktPH-mSc throughout the cell. The final technique used to display the LLSM data was through orthogonal planes. This generates 2D planes each 0.128 μm thick for the entire volume in xy, yz, and xz. Multiple methods were overlapped and shown side by side to effectively represent the data and labeled within each figure. It is important to note that all fluorescent information is retained within each individual pixel, providing a correlated visual increase with fluorescent probe localization.

## Visualization

The data visualization and analyses were performed using UCSF ChimeraX, developed by the Resource for Biocomputing, Visualization, and Informatics at the University of California, San Francisco, with support from National Institutes of Health R01-GM129325 and the Office of Cyber Infrastructure and Computational Biology, National Institute of Allergy and Infectious Diseases.

## LLSM

The Lattice Light Sheet Microscope referenced in this research was developed under license from Howard Hughes Medical Institute, Janelia Farms Research Campus (“Bessel Beam” patent applications 13/160,492 and 13/844,405).

## Author Contributions

S.E.Q. acquired and analyzed LLSM data and co-wrote the manuscript. L.H generated FLM cell lines used in experiments. J.G.K. Designed plasmid constructs and generated FLM cell lines. J.A.S. Performed SEM experiments and edited the manuscript. S.S. provided supervision and edited the manuscript. A.D.H. co-wrote the manuscript. R.B.A. built the LLSM and analysis cluster and provided supervision. N.W.T. designed experiments and co-wrote the manuscript. B.L.S. designed and performed initial experiments, assisted with data analysis and co-wrote the manuscript.

---

**Supplementary Figure 1.**
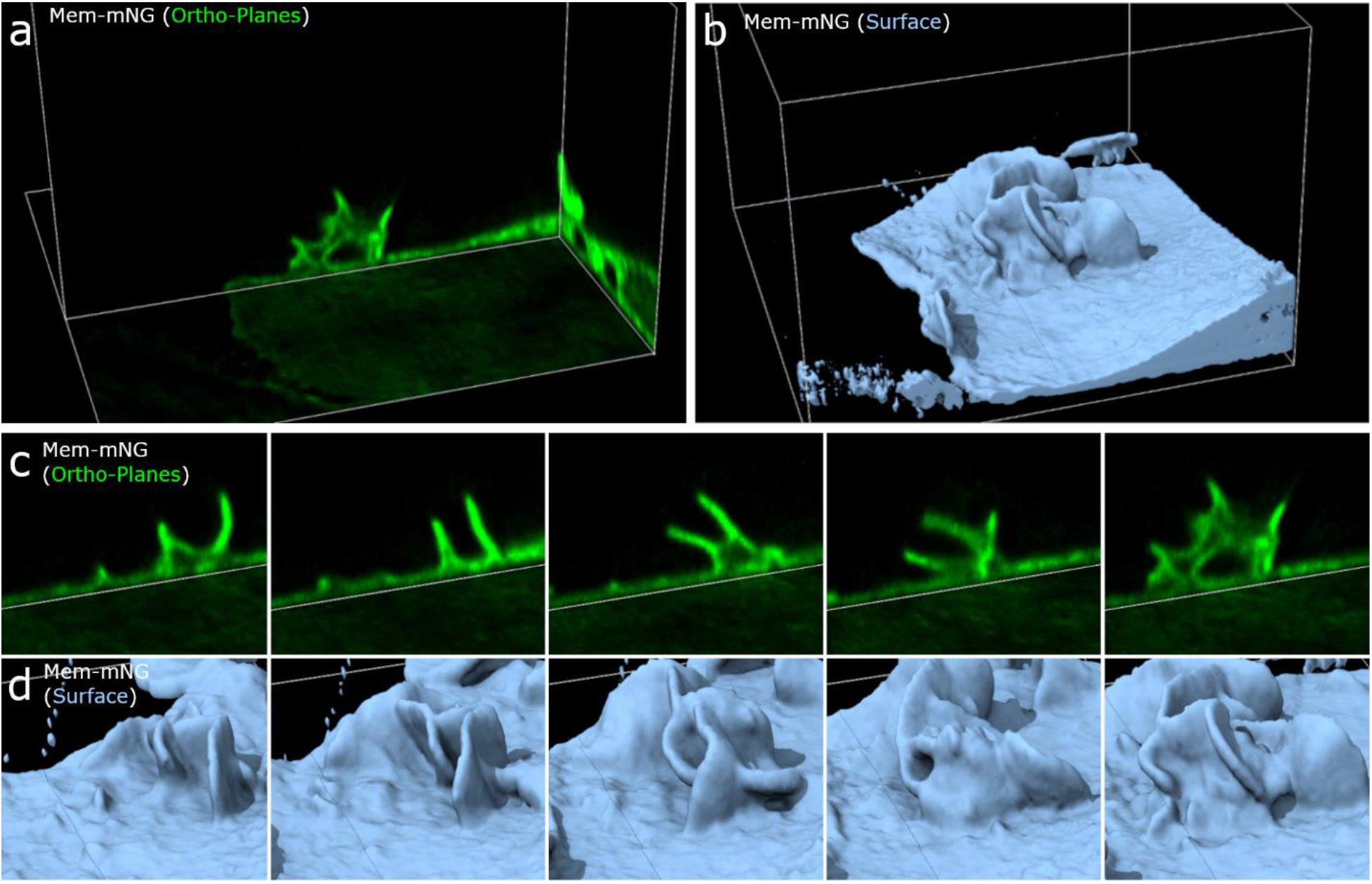
Constitutive macropinocytosis and the importance of reducing the dimensionality of data. a) Orthoplane (left) and isosurface (right) views of Mem-mNG show the subsurface macropinosome and the complex structure of the full surface (Region 20×20×15 μm). Orthoplane montage of Mem-mNG depicting constitutive planar view of macropinocytosis where two ruffles extend from the cell membrane, circularize, and connect to form a macropinosome. c) Close view of the respective orthoplane and isosurface views of Mem-mNG showing the complex structures and the multiple membrane ruffles involved in the macropinocytic event.

**Supplementary Figure 2.**
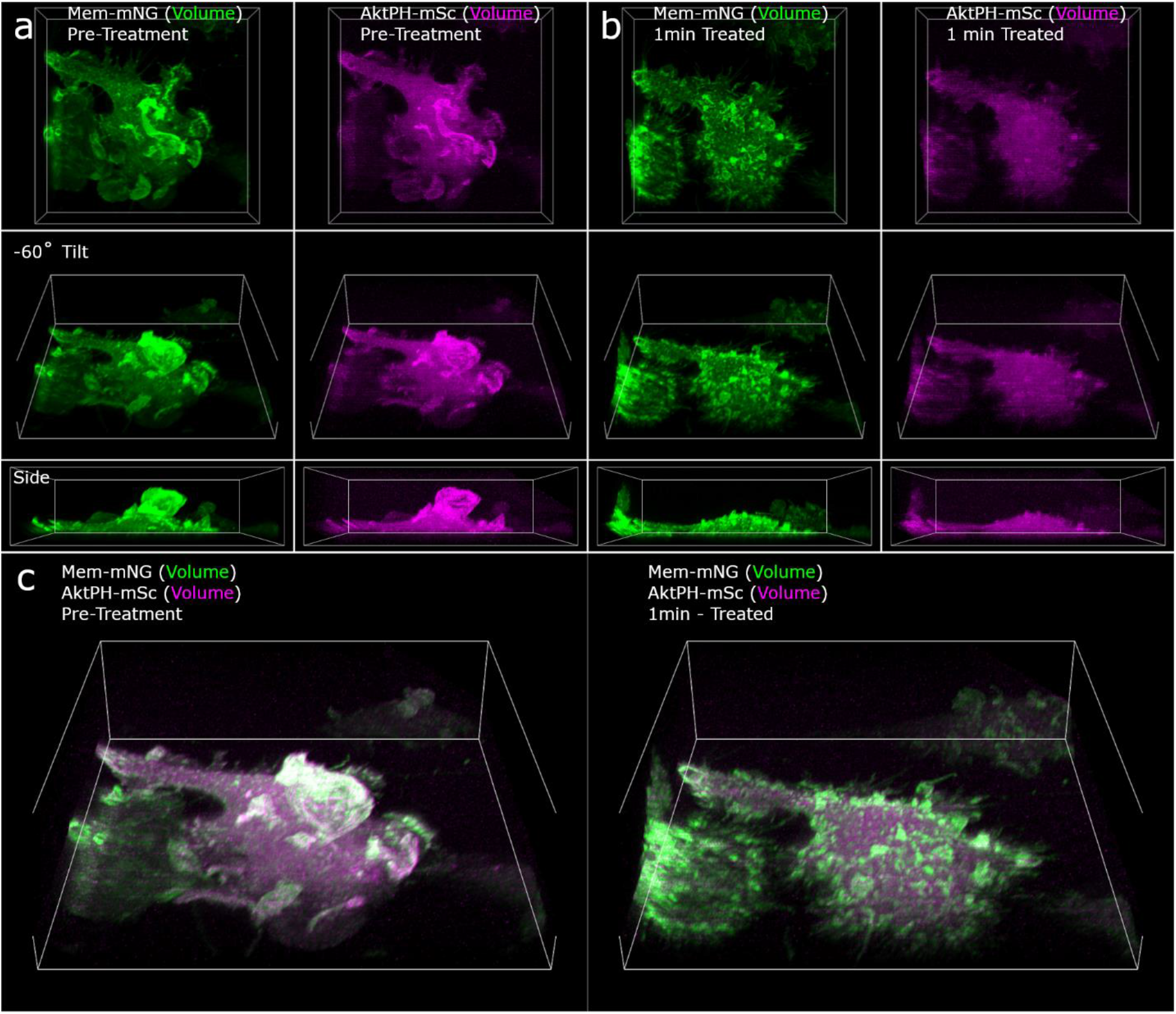
LY294002 drug treatment response. a) Two columns showing different angles of a split volume representations for Mem-mNG and AktPH-mSc of an untreated macrophage actively ruffling and creating macropinosomes (Region 71×69×20 μm). b) Two columns showing the same cell treated with LY294002 displayed as split volumes of Mem-mNG and AktPH-mSc one minute after drug treatment (Region 71×69×20 μm). c) Composite volume representations of pre-treatment (left) and one minute post LY294002 drug treatment (right) (Region 71×69×20 μm).

**Supplementary Figure 3.**
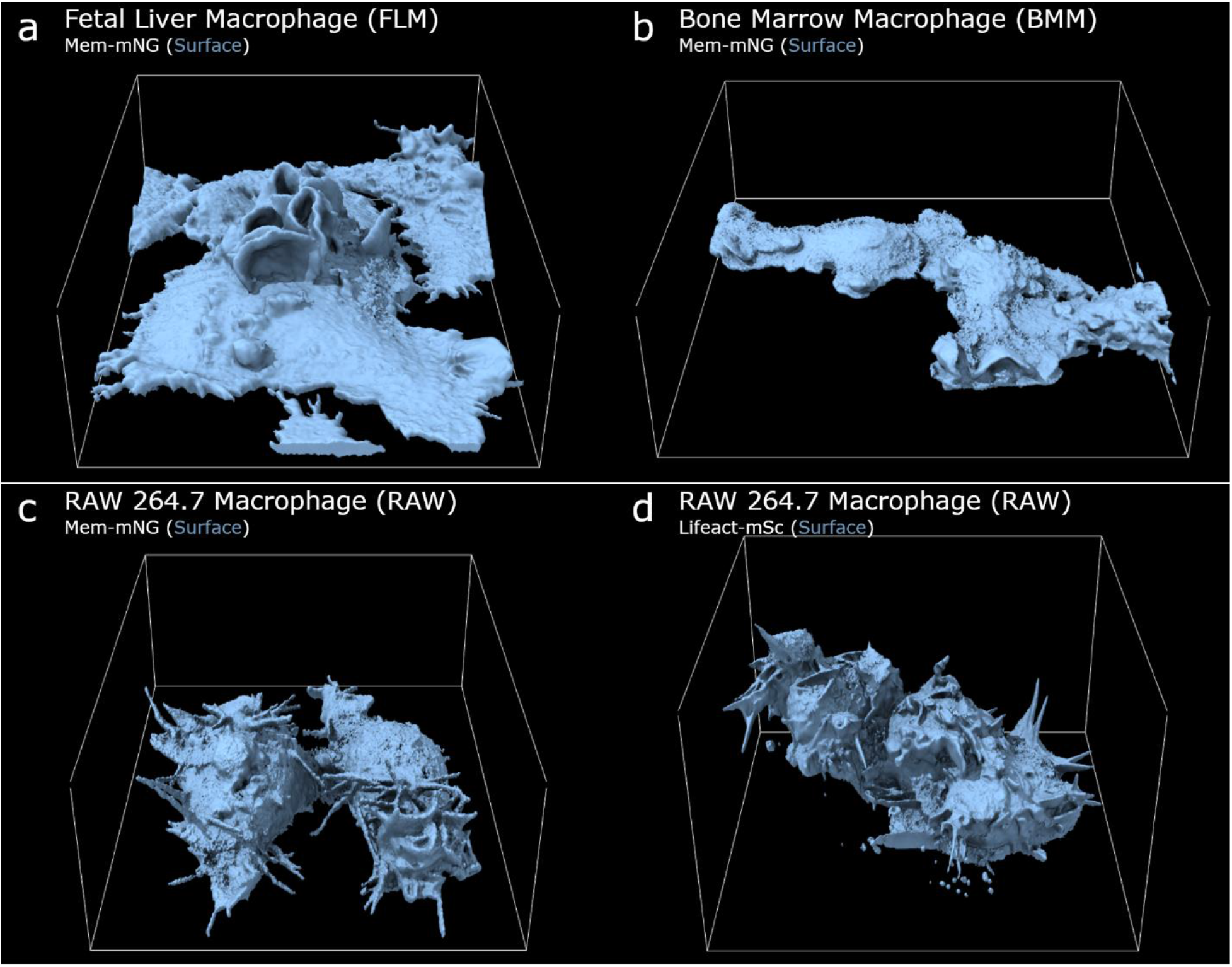
Cell line variability and cross comparison. Isosurface view of Mem-mNG showing the general membrane structure observed by LLSM: a) fetal liver macrophage (Region 47×59×19 μm), b) bone marrow derived macrophage (Region 71×68×22 μm), and c) RAW264.7 (Region 50×38×17 μm). d) Isosurface view of Lifeact-mSc expressed in RAW264.7 demonstrating a similar morphology with filopodial extensions (Region 49×49×30 μm).

## Supplementary Movie Legends

### Video 1. Corresponds to Fig. 1a, c

Isosurface and volumetric intensity renderings (Green Mem-mNG; Magenta AktPH-mSc) provides different methods of visualizing the formation (surface) and trafficking of macropinosomes and AktPH-mSc accumulation (volume). Movie timestamps highlight the following events: AktPH enriched ruffle development (02:42), newly formed macropinosomes (06:29; 08:51; 10:58), and post closure AktPH recruitment (11:19). Frame rate of ~7s and each region is 68×72×25 μm.

### Video 2. Corresponds to Fig. 1d

Isosurface rendering in conjunction with orthogonal planes moved through a volume provide ~0.1 μm thick planes to help visualize the internal and surface activity of Mem-mNG and AktPH-mSc including macropinosome closures (02:41; 05:15), membrane rich ruffles (00:00; 02:34; 05:08), previously formed internal macropinosomes (05:15 during scan), and mSc-Akt localization around a closed macropinosome (05:15 during scan). Framerate of ~7s and two subregions each 29×30×19 μm.

### Video 3. Corresponds to Fig. 2a

Isosurface and dual-volumetric intensity projection of Mem-mNG and AktPH-mSc with a 25-degree tilt shows a variety of formation events. Several early formations occur prior to relaxation of the plasma membrane (01:58), followed by the development of another AktPH rich ruffle (02:48), membrane closure into a macropinosome (03:07 -> 3:13), and finally post closure recruitment of AktPH (03:25). Several macropinosomes form that are indicated by the recruitment of AktPH post closure (3:57; 07:16) and subsequently trafficked toward one another to merge (4:53; 05:18; 05:30; 07:54). Frame rate of ~6.25s and region of 21×19×15um.

### Video 4. Corresponds to Fig. 2c

Mesh rendering of Mem-mNG and volumetric intensity projection of AktPH-mSc (Magenta-Hot) using a 90-degree tilt. The second play through contains a pause to emphasize the frame shown in Fig 2c and highlight the AktPH rich membrane ruffles (02:48) and the post closure recruitments (03:25; 03:57). Frame rate ~6.25s region of 21×19×15um.

### Video 5. Corresponds to Fig. 3a, b

Isosurface in conjunction with three volumetric intensity renderings (Green Mem-mNG; Magenta AktPH-mSc) display a traditional formation in an untreated cell including the initial ruffle (00:00) that vertically extends and begins to form a tidal wave (01:31) back toward the surface of the cell, with membrane scission (03:24 -> 03:31) and finally the post closure recruitment for the largest macropinosome (08:07). Frame rate of 7s and Region of 12×13×10 μm.

### Video 6. Corresponds to Fig. 3d, e

Isosurface with three volumetric intensity renderings (Green Mem-mNG; Magenta AktPH-mSc) on an LY294002 treated macrophage showing the initial ruffle with uniform AktPH throughout the cytosol and ruffle (00:00), attempted closure of the ruffle (00:30), continued compression of the attempted macropinosome (00:30 -> 02:27), becoming un-trackable within the cytosol with no AktPH recruitment to the attempted macropinosome. (Tracking done manually using orthogonal planes) Frame rate of 6.15s and Region 10×12×10 μm.

### Video 7. Corresponds to Supplementary Figure 2

Two color volumetric intensity renderings (Green Mem-mNG; Magenta AktPH-mSc) showing a pre-treated macrophage creating large dorsal ruffles rich with PI3K activity. After ~9 minutes of imaging, the cells were introduced to LY294002 and the same cell was reimaged showing a rapid reduction in ruffling and PI3K activity. After ~9 minutes of treatment, we show a second cell showing the start of returning ruffle activity on the same coverslip. Finally, a third cell was imaged starting at ~18 minutes into drug treatment showing the return of moderate ruffle formations but no increased PI3K activity (Region 71×69×20 μm).

### Video 8. Corresponds to Fig. 4a, b

Isosurface alongside three volumetric intensity renderings (Green Mem-mNG; Magenta AktPH-mSc) shows the formation of macropinosomes at the base of a larger ruffle. Membrane relaxes (00:56), small protrusions form with increased AktPH (01:17), two small ruffles, one in the back and one in the front mergers with the larger ruffle (01:32 -> 01:39) followed by post closure AktPH recruitment (01:53). Frame rate of 7s and region of 11×9×12 μm.

### Video 9. Corresponds to Fig. 5c

Isosurface and dual-volumetric intensity projection of Mem-mNG and AktPH-mSc showing a smooth and relaxed membrane (00:00). A single macropinosome forms (05:06) followed by a large increased in membrane activity (06:35) resulting in a significant number of macropinosomes, indicated by the post closure AktPH recruitment, that turns into the chaotic membrane structure (06:35 -> 13:18). Frame rate of 8s and region size of 27×22×16 μm.

### Video 10. Corresponds to Fig. 5e

Side view of Mem-mNG isosurface and mesh membrane with volumetric AktPH (Magenta Hot) shows the continued AktPH localization within the extending membrane structure (06:35). Utilizing the Magenta-Hot LUT regions displaying in white represent the increase in AktPH post macropinosome closure signifying a formed macropinosome (05:55; 06:59; 08:12; 11:50). Frame rate of 8s and region size of 27×22×16 μm.

### Video 11. Corresponds to Fig. 6a, b, c

CSF-1 starved macrophage displayed using Mem-mNG isosurface/mesh/volume and AktPH volumes as magenta-hot under the mesh and Magenta alongside the volume membrane. The macrophage was imaged 07:41 prior to stimulation and reimaged one minute after CSF-1 stimulation (08:41) providing time to ensure instrument and imaging conditions had not changed. The starved cell ruffled and formed macropinosomes similar to the conventional cells (00:06; 01:46; 03:26) and upon stimulation (08:41) a large circular dorsal ruffle forms corralling the AktPH to one concentrated spot in the cell (16:22). Frame rate of 6.25s and region size of 49×60 μm.

### Video 12. Corresponds to Fig. 6d

The brightfield view starts promptly after stimulation showing the majority of macrophages performing the similar dorsal membrane clearing seen in the LLSM imaging.

### Video 13. Corresponds to Fig. 7a, b. LPS Stimulation

Isosurface of Mem-mNG and dual-color volumetric intensity projections of Mem-mNG and AktPH-mSc showing the activity of an LPS treated cell. Initial imaging starts (00:00) with a cluster of membrane rich in AktPH that goes on to create many macropinosomes as it expands toward the upper right region of the field of view. The activity changes directions toward the upper left region of the cell and proceeds to move counterclockwise (01:14), over the nucleus and back to the initial location ending at (04:42). Additional macropinosomes are seen forming on the left region with the increased AktPH flare up post closure (04:42). Finally, several formations occur in the bottom right of the cell (05:05 – 08:18) many of which go on to merge with one another. Framerate of ~7s and Region of 68×77×21 μm.

### Video 14. Corresponds to Fig. 7c, d. Non-treated control

Isosurface of Mem-mNG and dual-color volumetric intensity projections of Mem-mNG and AktPH-mSc showing the imaging of a nontreated cell (10:47). Two AktPH rich regions of membrane ruffling are seen in the bottom right of the cell (02:17) that form several small macropinosomes, indicated by a spike in AktPH around the formed macropinosome. The cell shows activity that is representative of the untreated experiments including macropinosome formations, exploration, and overall membrane ruffling. Framerate of ~6.5s and Region of 68×77×21 μm.

### Video 15. Corresponds to Supplementary Figure 3b

Isosurface rendering of Mem-mNG in a BMM. The cell shows activity that is representative of the BMM population including macropinosome formations, surface spreading, and overall membrane ruffling. Framerate of ~4.2s and region 71×68×22 μm).

### Video 16. Corresponds to Supplementary Figure 3c

Isosurface rendering of Mem-mNG in an RAW264.7 cell. The cell shows activity that is representative of the RAW264.7 population including tentpole extensions, rounded shape, and general membrane activity. Framerate of ~3.5s and region 50×38×17 μm).

## References

1 Freeman, S. A. et al. Lipid-gated monovalent ion fluxes regulate endocytic traffic and support immune surveillance. Science 367, 301–305, doi:10.1126/science.aaw9544 (2020).

2 Bohdanowicz, M. & Grinstein, S. Role of phospholipids in endocytosis, phagocytosis, and macropinocytosis. Physiol Rev 93, 69–106, doi:10.1152/physrev.00002.2012 (2013).

3 Doodnauth, S. A., Grinstein, S. & Maxson, M. E. Constitutive and stimulated macropinocytosis in macrophages: roles in immunity and in the pathogenesis of atherosclerosis. Philosophical Transactions of the Royal Society B: Biological Sciences 374, 20180147, doi:doi:10.1098/rstb.2018.0147 (2019).

4 Kerr, M. C. & Teasdale, R. D. Defining Macropinocytosis. Traffic 10, 364–371, doi:10.1111/j.1600-0854.2009.00878.x (2009).

5 Redka, D. y. S., Gütschow, M., Grinstein, S. & Canton, J. Differential ability of proinflammatory and anti-inflammatory macrophages to perform macropinocytosis. Molecular Biology of the Cell 29, 53–65, doi:10.1091/mbc.E17-06-0419 (2018).

6 Egami, Y., Taguchi, T., Maekawa, M., Arai, H. & Araki, N. Small GTPases and phosphoinositides in the regulatory mechanisms of macropinosome formation and maturation. Frontiers in Physiology 5, doi:10.3389/fphys.2014.00374 (2014).

7 Swanson, J. A. & Yoshida, S. Macropinosomes as units of signal transduction. Philosophical Transactions of the Royal Society B: Biological Sciences 374, 20180157, doi:doi:10.1098/rstb.2018.0157 (2019).

8 Yoshida, S. et al. Differential signaling during macropinocytosis in response to M-CSF and PMA in macrophages. Frontiers in physiology 6, 8–8, doi:10.3389/fphys.2015.00008 (2015).

9 Lou, J., Low-Nam, S., Kerkvliet, J. & Hoppe, A. Delivery of CSF-1R to the lumen of macropinosomes promotes its destruction in macrophages. Journal of cell science 127, doi:10.1242/jcs.154393 (2014).

10 Bloomfield, G. & Kay, R. R. Uses and abuses of macropinocytosis. Journal of Cell Science 129, 2697, doi:10.1242/jcs.176149 (2016).

11 Pacitto, R., Gaeta, I., Swanson, J. A. & Yoshida, S. CXCL12-induced macropinocytosis modulates two distinct pathways to activate mTORC1 in macrophages. Journal of Leukocyte Biology 101, 683–692, doi:https://doi.org/10.1189/jlb.2A0316-141RR (2017).

12 Condon, N. D. et al. Macropinosome formation by tent pole ruffling in macrophages. Journal of Cell Biology 217, 3873–3885, doi:10.1083/jcb.201804137 (2018).

13 Swanson, J. A. Shaping cups into phagosomes and macropinosomes. Nature Reviews Molecular Cell Biology 9, 639–649, doi:10.1038/nrm2447 (2008).

14 Botelho, R. J. et al. Localized Biphasic Changes in Phosphatidylinositol-4,5-Bisphosphate at Sites of Phagocytosis. Journal of Cell Biology 151, 1353–1368, doi:10.1083/jcb.151.7.1353 (2000).

15 Freeman, S. A. & Grinstein, S. Phagocytosis: receptors, signal integration, and the cytoskeleton. Immunol Rev 262, 193–215, doi:10.1111/imr.12212 (2014).

16 King, J. S. & Kay, R. R. The origins and evolution of macropinocytosis. Philosophical Transactions of the Royal Society B: Biological Sciences 374, 20180158, doi:doi:10.1098/rstb.2018.0158 (2019).

17 Yoshida, S., Hoppe, A. D., Araki, N. & Swanson, J. A. Sequential signaling in plasma-membrane domains during macropinosome formation in macrophages. Journal of Cell Science 122, 3250–3261, doi:10.1242/jcs.053207 (2009).

18 Veltman, D. M. et al. A plasma membrane template for macropinocytic cups. eLife 5, e20085, doi:10.7554/eLife.20085 (2016).

19 Donaldson, J. G. Macropinosome formation, maturation and membrane recycling: lessons from clathrin-independent endosomal membrane systems. Philosophical Transactions of the Royal Society B: Biological Sciences 374, 20180148, doi:doi:10.1098/rstb.2018.0148 (2019).

20 Maekawa, M. et al. Sequential breakdown of 3-phosphorylated phosphoinositides is essential for the completion of macropinocytosis. Proceedings of the National Academy of Sciences 111, E978, doi:10.1073/pnas.1311029111 (2014).

21 Buckley, C. M. & King, J. S. Drinking problems: mechanisms of macropinosome formation and maturation. The FEBS Journal 284, 3778–3790, doi:10.1111/febs.14115 (2017).

22 Araki, N., Egami, Y., Watanabe, Y. & Hatae, T. Phosphoinositide metabolism during membrane ruffling and macropinosome formation in EGF-stimulated A431 cells. Exp Cell Res 313, 1496–1507, doi:10.1016/j.yexcr.2007.02.012 (2007).

23 Welliver, T. P. & Swanson, J. A. A growth factor signaling cascade confined to circular ruffles in macrophages. Biology Open 1, 754, doi:10.1242/bio.20121784 (2012).

24 Chamberlain, L. M., Godek, M. L., Gonzalez-Juarrero, M. & Grainger, D. W. Phenotypic non-equivalence of murine (monocyte-) macrophage cells in biomaterial and inflammatory models. Journal of Biomedical Materials Research Part A 88A, 858–871, doi:https://doi.org/10.1002/jbm.a.31930 (2009).

25 Gao, L., Shao, L., Chen, B.-C. & Betzig, E. 3D live fluorescence imaging of cellular dynamics using Bessel beam plane illumination microscopy. Nature Protocols 9, 1083–1101, doi:10.1038/nprot.2014.087 (2014).

26 Goddard, T. D. et al. UCSF ChimeraX: Meeting modern challenges in visualization and analysis. Protein Sci 27, 14–25, doi:10.1002/pro.3235 (2018).

27 Araki, N., Johnson, M. T. & Swanson, J. A. A role for phosphoinositide 3-kinase in the completion of macropinocytosis and phagocytosis by macrophages. Journal of Cell Biology 135, 1249–1260, doi:10.1083/jcb.135.5.1249 (1996).

28 Luyendyk, J. P. et al. Genetic Analysis of the Role of the PI3K-Akt Pathway in Lipopolysaccharide-Induced Cytokine and Tissue Factor Gene Expression in Monocytes/Macrophages. The Journal of Immunology 180, 4218–4226, doi:10.4049/jimmunol.180.6.4218 (2008).

29 Hoeller, O. et al. Two distinct functions for PI3-kinases in macropinocytosis. Journal of Cell Science 126, 4296, doi:10.1242/jcs.134015 (2013).

30 Hacker, U., Albrecht, R. & Maniak, M. Fluid-phase uptake by macropinocytosis in Dictyostelium. Journal of Cell Science 110, 105–112, doi:10.1242/jcs.110.2.105 (1997).

31 Williams, T. D., Peak-Chew, S.-Y., Paschke, P. & Kay, R. R. Akt and SGK protein kinases are required for efficient feeding by macropinocytosis. Journal of Cell Science 132, doi:10.1242/jcs.224998 (2019).

32 Stewart, S. A. et al. Lentivirus-delivered stable gene silencing by RNAi in primary cells. Rna 9, 493–501, doi:10.1261/rna.2192803 (2003).

33 Sancak, Y. et al. The Rag GTPases bind raptor and mediate amino acid signaling to mTORC1. Science 320, 1496–1501, doi:10.1126/science.1157535 (2008).

34 Chertkova, A. O. et al. Robust and Bright Genetically Encoded Fluorescent Markers for Highlighting Structures and Compartments in Mammalian Cells. bioRxiv, 160374, doi:10.1101/160374 (2020).

35 Fejer, G. et al. Nontransformed, GM-CSF–dependent macrophage lines are a unique model to study tissue macrophage functions. Proceedings of the National Academy of Sciences 110, E2191–E2198, doi:10.1073/pnas.1302877110 (2013).

36 Lebedev, M., Swaminathan, P., Kerkvlie, J. G., Hoppe, A. D. & Thiex, N. Immortal fetal liver macrophages as a new model for studying macrophage function. Molecular Biology of the Cell 27(2016).

37 Stanley, E. R., Cifone, M., Heard, P. M. & Defendi, V. Factors regulating macrophage production and growth: identity of colony-stimulating factor and macrophage growth factor. J Exp Med 143, 631–647, doi:10.1084/jem.143.3.631 (1976).

38 Waheed, A. & Shadduck, R. K. Purification and properties of L cell-derived colony-stimulating factor. J Lab Clin Med 94, 180–193 (1979).

39 tlambert03/LLSpy: v0.4.8 v. 0.4.8 (Zenodo, 2019).

